# The Reactobiome Unravels a New Paradigm in Human Gut Microbiome Metabolism

**DOI:** 10.1101/2021.02.01.428114

**Authors:** Gholamreza Bidkhori, Sunjae Lee, Lindsey A. Edwards, Emmanuelle Le Chatelier, Mathieu Almeida, Bouchra Ezzamouri, Florian Plaza Onate, Nicolas Ponte, Debbie L. Shawcross, Gordon Proctor, Lars Nielsen, Jens Nielsen, Mathias Uhlen, Stanislav Dusko Ehrlich, Saeed Shoaie

**Affiliations:** Centre for Host-Microbiome Interactions, Faculty of Dentistry, Oral & Craniofacial Sciences, King’s College London, SE1 9RT, UK; Liver Sciences, Department of Inflammation Biology, School of Immunology and Microbial Sciences, King’s College London, London, UK; INRAE, Institut National de la Recherche Agronomique, US1367 MetaGenoPolis, 78350, Jouy en Josas, France; Novo Nordisk Foundation Center for Biosustainability, Technical University of Denmark, DK-2800 Kgs. Lyngby, Denmark; Department of Biology and Biological Engineering, Kemivägen 10, Chalmers University of Technology, SE-412 96, Gothenburg, Sweden; Science for Life Laboratory, KTH - Royal Institute of Technology, 171 21, Stockholm, Sweden; AIVIVO Ltd. Unit 25, Bio-innovation centre, Cambridge Science park, Cambridge, UK

## Abstract

Changes in microbial metabolism have been used as the main approach to assess function and elucidate environmental and host-microbiome interactions. This can be hampered by uncharacterised metagenome species and lack of metabolic annotation. To address this, we present a comprehensive computational platform for population stratification based on microbiome composition, the underlying metabolic potential and generation of metagenome species and community level metabolic models. We revisit the concepts of enterotype and microbiome richness introducing the reactobiome as a stratification method to unravel the metabolic features of the human gut microbiome. The reactobiome encapsulates resilience and microbiome dysbiosis at a functional level. We describe five reactotypes in healthy populations from 16 countries, with specific amino acid, carbohydrate and xenobiotic metabolic features. The validity of the approach was tested to unravel host-microbiome and environmental interactions by applying the reactobiome analysis on a one-year Swedish longitudinal cohort, integrating gut metagenomics, plasma metabolomics and clinical data.

## Introduction

Utilising enumeration of metagenomics to determine microbial composition such as enterotype^1,2^ and richness^3^ can only partially explain the microbiome drivers of health and chronic disease. Individual and population stratifications can be greatly complemented by quantitative integration with host physiology and therefore provide a mechanistic causality in host-microbiome interactions^4^. Metabolism is one of the main underlying mechanisms that shapes the resilience of the microbiome and determines host homeostasis; quantitative measurements of faecal and blood metabolites and analysis of their associations with gut microbial composition, has proven to be a fruitful approach to explore phenotypes and characterize the interplay of bacterial diversity with local and circulating metabolites^5,6^.

Constraint-based modelling, using GEnome-scale Metabolic models (GEMs) can provide a mechanistic understanding of the microbial metabolism and of genotype-phenotype relationships in host-microbiome interactions ^7,8^. GEMs have been applied to simple microbial community modelling in the gut and demonstrated its power to identify the key metabolic contribution of bacteria in the ecosystem and to host metabolism^9-11^. Previously, a number of human gut bacterial GEMs have been generated, mainly based on the available whole genome sequence, and used to investigate the bacterial growth rates under different nutrient availabilities on a community level ^9,10,12-15^. However, the challenge remains to be able to use metagenomic data directly to reconstruct the GEMs for microbiome uncultured genomes and apply the models to personalized microbiome investigation, or to identifying individuals endowed with different types of microbial metabolism, by unsupervised clustering. Using the latest developments of integrated non-redundant microbial gene catalogues^16,17^ and metagenome binning^18,19^ to acquire more genomes, could be a comprehensive approach to obtain high-quality GEMs.

Here, we present the toolbox for <u>MI</u>crobial and personalized <u>G</u>EM, <u>RE</u>actobiome and community NEtwork modelling (MIGRENE), which enables generating species and community-level models to be applied to personalized microbiome studies. Using this toolbox, we developed a new paradigm, the ‘reactobiome’, revealing novel aspects of the function of the gut microbiome. We propose a new microbiome classification, termed ‘reactotype’, and identify five different reactotypes. We used the reactobiome to analyse the association of the gut microbiome over 1-year with matched plasma biochemistry and metabolome sampling^20^, at four-time points, to give a detailed picture encapsulating the complexity of the host-microbiome interactions.

## Results

### Characteristics of the MAGMA and reactobiome in the global gut microbiome of healthy individuals

We developed the MIGRENE toolbox to investigate the microbiome using metagenome species with integrated microbial gene catalogues (Fig. 1a, Extended Data Fig. 1a-c). The toolbox generates high-quality GEMs; here they are based on the gut microbial gene catalogue^21^ and metagenomic species pan-genomes (MSPs)^18^, and are referred to as <u>M</u>etagenome species <u>A</u>ssembled <u>G</u>enome-scale <u>M</u>et<u>A</u>bolic model (MAGMA). From the 1898 gut MSPs, 1706 bacterial MSPs were taxonomically classified and used to reconstruct MAGMAs. We successfully generated 1333 functional MAGMAs (Supplementary Table 1), with completeness 83.6±22.5%. The completeness of the 373 non-functional models was 48.4±35.6% (Extended Data Fig. 2a); future completion of genomes of these species will allow generation of the corresponding functional models. Functionality of the models was assessed by constraint-based modelling, considering biomass as the objective function and western diet macronutrients as the main substrate (Extended Data Fig. 2b). The current MAGMAs contain a median number of 1478 reactions and average number of 647 genes from an average of 1545 annotated genes in MSPs (Extended Data Fig. 2c). Expectedly, the number of reactions in MAGMAs and genes in MSPs were correlated (adj. R^2^ = 0.61, Extended Data Fig. 2d). Median gap filling was only 4.13%; 57% of the added reactions were from the highest taxonomy-level (760 genus and 350 family) that could be determined (Extended Data Fig. 2e-g). Compared to other publicly available GEMs resources ^10,22^, MAGMAs had superior connectivity (Extended Data Fig. 2h-j, Supplementary Table 2). Moreover, MAGMAs reactions clustered by phylum and family level taxonomic classification (Extended Data Fig. 2k-n).

**Fig 1.**
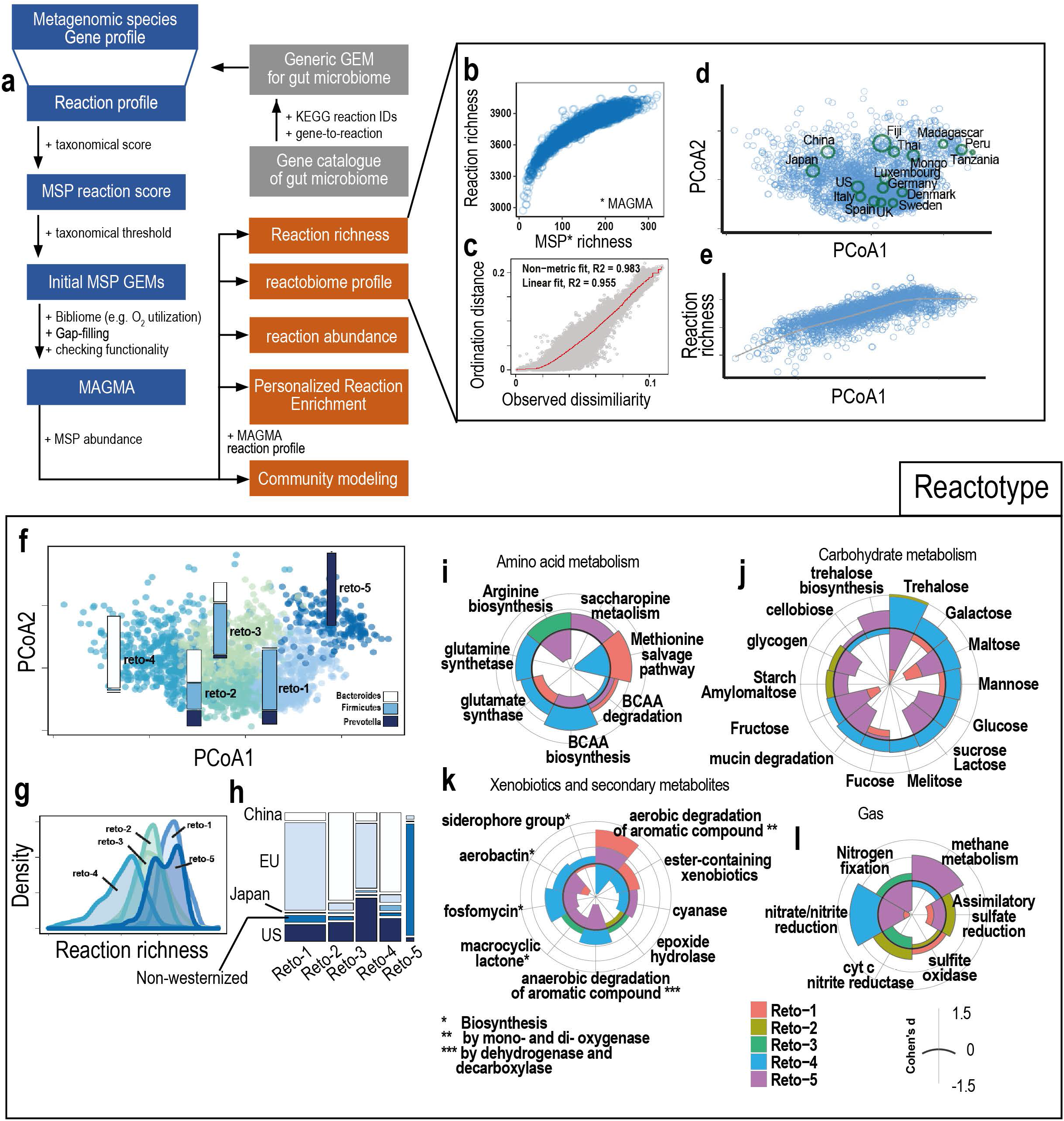
The MIGRENE toolbox, global landscape of reaction richness and the reactobiome in healthy samples and characterization of the reactotypes. **a**, the workflow of the MIGRENE toolbox generates utilize any non-redundant microbial gene catalogue, metagenome species to generate a microbiome generic metabolic and species-specific model (<u>M</u>SP <u>A</u>ssociated <u>G</u>enome scale <u>M</u>et<u>A</u>bolic models; MAGMA). The MIGRENE toolbox can determine personalized metabolic microbiome profile and community metabolic modelling. (Detailed in Extended Data Fig. 1). **b**, scatter plot of the MSP richness, that MAGMA could be generated (1333 MSPs), and reaction richness in healthy gut samples (n=2911). **c**, NMDS Shepard plot shows the relationship between the actual dissimilarities (x-axis) between samples and the ordination distances (y-axis) by reactobiome. The fit between dissimilarity and ordination distances is indicated by red line. **d**, visualization of 2911 samples with Principal Coordinates Analysis (PCoA) using reactobiome; projected centroid of each country from the PCoA is shown as circle, the size indicates the average standard deviation of PCoA1 and PCoA2. **e**, scatter plot for reaction richness and the first dimension of the PCoA plot. **f**, visualization of the reactotypes (reto-1 to reto-5) on the PCoA plot; the bars show the proportion of enterotypes in each reactotype. **g**, the density plot of reaction richness for the reactotypes. **h**, the proportion of geographical regions in reactotypes. **i-l**, radar plots showing the fraction of functional terms in reto1 to reto 5, tested by enriched/depleted reactions of the reactobiome based on Cohens’ d effect sizes of Wilcoxon one-sided tests (positive and negative Cohens’ d respectively indicate enriched and depleted reactions).

Recently, we developed the human gut microbiome atlas (HGMA; www.microbiomeatlas.com) to investigate the global characterization of the human gut microbiome in health and disease. From the HGMA, using MSP profile of 2911 healthy samples from 16 countries together with the MAGMA reaction pool we generated community modelling, personalized reaction abundance, enrichment and richness, and the reactobiome (an aggregate of the metabolic repertoires of an individual gut microbiome) using the MIGRENE toolbox (Extended Data Fig. 1d, Supplementary Tables 3-6). For the samples from healthy subjects, reaction richness exponentially increased with the number of MSPs and gene richness (Fig. 1b, Extended Data Fig. 2o-p, Supplementary Table 4). The reactobiome profile substantially improved the fit between the ordination distances and observed dissimilarities in two dimensions compared to the MSP abundance (Fig. 1c, Extended Data Fig. 2q, Supplementary Table 5, Method). Additionally, the scree plot showed a clear elbow at second dimension for the reactobiome, indicating that 2 dimensions are significant; in contrast, two-dimensional reduction space was inadequate for MSP abundances (Fig. 1d, Extended Data Fig. 2r-s). Reaction richness projection on the reactobiome in samples from healthy subjects gradually increased along the first dimension of PCoA plot. Projected centroid of each country from the PCoA revealed geographical differences in the gut reactobiome, Asian countries (China and Japan), US and Europe, and non-westernized countries grouping together, along the increasing reaction richness gradient (Fig. 1d-e).

### Reactotypes; stratifying individual gut microbiome based on the reactobiome

To stratify the biochemical state of an individual gut microbiome, i.e. reactobiome, we performed unsupervised clustering, Dirichlet multinomial mixture (DMM)^23^, and calculated the number of optimal clusters based on fitting Dirichlet mixtures by minimising the negative log posterior and then Laplace approximation (Extended Data Fig. 2t, Method). We identified five different “Reactotypes” (reto-1 to reto-5) (Supplementary Table 7) differing by enterotype distribution (Fig. 1f, Method): reto-1 and -3 were enriched in the Firmicutes enterotype, while reto -4 and -5 were enriched in the Bacteroidetes and Prevotella enterotypes, respectively. The five reactotypes also differed in reaction and MAGMA richness (Fig. 1g, Extended Fig 3a) and in geographical distribution (Fig. 1h). Reto-1 and -5 had high richness, reto-2 and -3 intermediate and reto-4 had low richness; reto-1 and -3 were predominant in westernized countries (EU and US), reto-2 and reto-4 in China and reto-5 was almost exclusively present in non-westernized countries. Interestingly, the reactotypes were clustered based on their associated metadata (Extended Data Fig. 3b). To further test whether the reactotypes differ significantly we used supervised machine learning models and cross validation (Method). The classification accuracy of the prediction models ranged from 0.98 to 1 (Extended Data Fig. 3c-d).

We further defined key features of each reactotype by non-parametric tests (Kruskal-Wallis test and Dunn’s multiple comparisons; Supplementary Table 8 and Method). To find the reactotype-specific enriched and depleted reactions, the Cohen’s d effect size of one-sided Wilcoxon test results was performed (|Cohen’s d| >= 0.5, Methods) (Supplementary Table 9). Reto-1, reto-3 and reto-5 showed a greater number of enriched and depleted reactions compared to reto-2 and reto-3 (Extended Data Fig. 3e). Arginine biosynthesis was enriched in reto-3, and reto-5 was dominant in saccharopine and lysine metabolism. Methionine biosynthesis was enriched in reto-1. Reto-4 was dominant for producing the branched chain amino acids (BCAA), glutamate and glutamine, while the BCAA degradation was enriched in reto-1 and 5 (Fig. 1i-l, Extended Data Fig. 3f). BCAA levels, which reflect the synthesis/degradation balance, are correlated with insulin resistance ^24^; they are expected to be high in low richness reto-4 and low in high-richness reto-1 and -5, consistent with the low microbiome richness being a risk factor for type 2 diabetes and other metabolic syndrome related chronic diseases^3^. Carbohydrate metabolism, mono and di-saccharide degradation and mucin degradation were dominant in reto-4 and lowest in reto-1 and 5; simple sugar utilisation may reflect a sucrose-enriched diet, also conducive to metabolic syndrome related chronic diseases. Consistently, starch degradation (alpha amylase) was low in reto-1 and reto-5 and high in reto-2 and reto-4 (Extended Data Fig. 4), while cellobiose degradation was high in reto-5; this may reflect a starch-enriched diet in low richness reactobiomes and fibre-enriched diet, reported to counter loss of microbiome richness and improve the insulin resistance ^25^, in high richness reto-5. Glycogen, starch and amylomaltose degradation were the key features in the reto-2, while glycogen end-products appear to be different between the reactotypes, e.g. reto-1 and 5 had the highest production of 1,5-Anhydroglucitol (1,5-AG) while 1,5-AnhydroD-mannitol production was dominant in reto-4 (Extended Data Fig. 4). We also observed reto-1 and 5 were enriched in xenobiotic degradation, while reto-3 was a macrocyclic lactone producer and reto-4 was secondary metabolism enriched for biosynthesis of aerobactin, fosfomycin and sidrophores. Reto-1, 4 and 5 had the highest capability for aromatic compound degradation. Reto-1, similar to reto-5, was dominant in monooxygenase and dioxygenase activity while reto-4 was dominant in anaerobic degradation of aromatic compounds. Reto-1 had the highest capacity for detoxifying ester-containing xenobiotics. Reto-3 had enriched xenobiotic degradation by epoxidase hydrolase and reto-5 was cyanase enriched. We also found reto-2 was dominant in H_2_S production, reto-3 in nitrogen fixation, reto-4 in NH_4_^+^ production and reto-5 in methane production. Reto-4 had the highest capacity to produce branched chain fatty acids (BCFAs), and was enriched in fatty acid metabolism, amino sugars and mucin degradation.

### Mucin degradation increased leucine, isoleucine and proline production in the low richness reactotype

As reaction richness appears to be one of the main dictators of the reactobiome, we focused on the comparison of the high richness reto-1 and the low richness reto-4, dominant in EU and China and denoted EHR (European high richness) and CLR (Chinese low richness) reactotype, respectively. Some 1735 reactions had significantly different abundances between the two (Wilcoxon signed-rank test, q-value <1×10^−5^, Supplementary Table 10). Among these, 17 mucin degrading reactions were significantly enriched in CLR: sialidase-1 (NEU1, neuraminidase 1), beta-galactosidase, alpha-fucosidase and glucosylceramidase (Fig. 2a, Supplementary Table 11). Coherently, the reactions involved in metabolism of amino sugars and monosaccharides, the metabolites that can result from mucin degradation, were also enriched in CLR samples (Extended Data Fig. 5a, Supplementary Fig. 1 and S2). Here we observed, decreased availability of free mannose in reto-4 in comparison to reto-1 and reto-5, due to enhanced consumption, which has the potential to enhance pathogenic *E. coli* adherence to mucosal epithelial barriers ^26^. We also observed the same trend between reto-4 and reto-5 (Supplementary Table 12).

**Fig 2.**
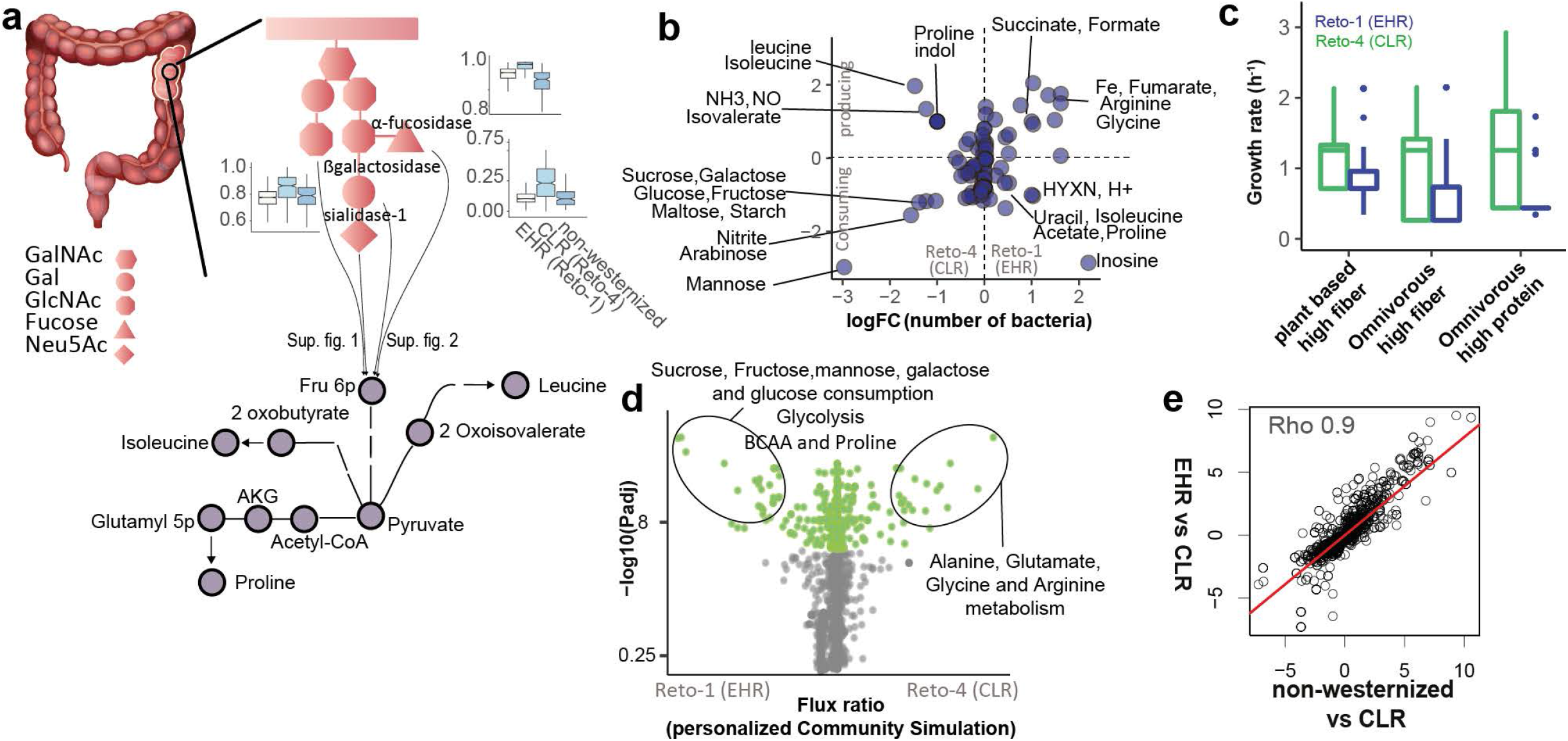
Characterization of personalized gut microbiome metabolism in European high richness (EHR, reto-1) and Chinese low richness (CLR, reto-4), reactotypes. **a**, enriched metabolic connections of host-microbiota in the CLR (reto-4). General structure of host-secreted mucin 2 glycoproteins (green)^51^ degraded by highly abundant carbohydrate-active enzymes in reto-4 compared to HER and non-westernized regions. The sugar source from mucin degradation or diet leads to increased production of. leucine, isoleucine and proline, in CLR. **b**, log fold change of the production/consumption flux for EHR vs CLR (y-axis) and the log fold change of the top 50-ranked MSPs in EHR and CLR (x-axis) producing/consuming the corresponding metabolites. **c**, the growth rate of the bacteria in 3 different diets (plant- and omnivore-based diet). **d**, the differences of average flux ratio between personalized gut bacterial community of EHR vs CLR. **e**, similarity of differences between non-westernized (reto-5) vs CLR and EHR vs CLR. The log fold change of variables significantly different between EHR vs CLR paired with the variables significantly different between non-westernized vs CLR (Spearman’s *r* = 0.9).

**Fig 3.**
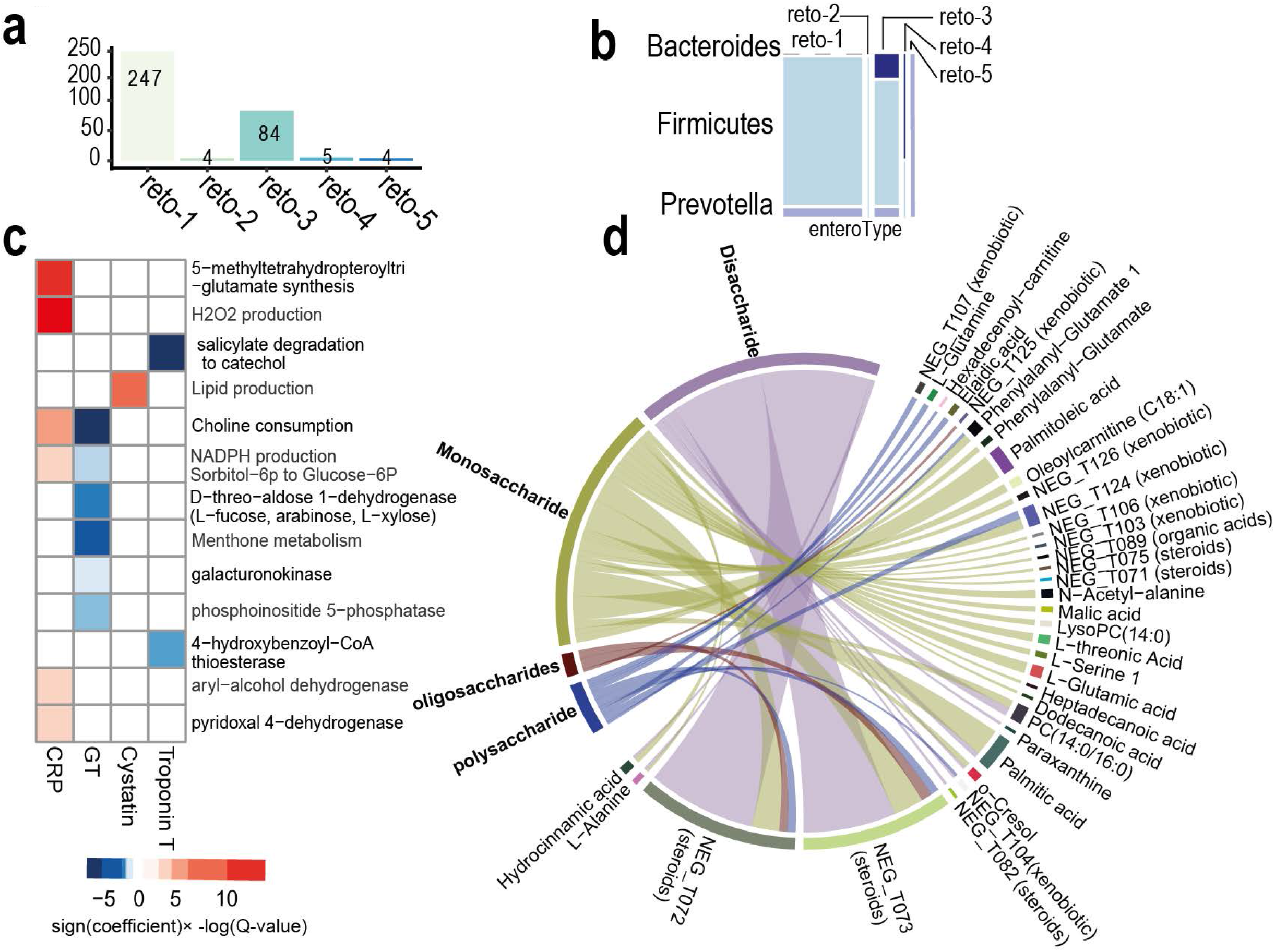
Interplay of host clinical parameters and plasma metabolites with variation of gut reactobiome. **a**, numbers of samples of each reactotype in the Swedish wellness cohort. **b**, the proportion of enterotypes for wellness reactotypes. The thickness of the bars shows the number of samples for each reactotype. **c**, the heatmap indicates the association between clinical biochemical factors and variations of the gut reactobiome at FDR < 0.01, using a multivariate random effects model. **d**, chord diagram showing the association between reactions of the reactobiome involved in consumption of carbohydrates with plasma metabolites at FDR 0.01, using multivariate random effects model.

The metabolic capacity of EHR and CLR was investigated using constrained-based modelling of the MAGMAs from the top 50-ranked MSPs significantly abundant in one or the other (Wilcoxon test, q-value < 0.05, Supplementary Table 13). Simulations were based on the maximization of the biomass as the objective function and high protein omnivorous diet as the main substrate (Method). The simulations showed amino acid production/consumption was different between CLR and EHR (Fisher’s exact test, FDR <0.01, Fig. 2b). Three amino acids, leucine, isoleucine and proline were produced in CLR and consumed in EHR (Supplementary Table 14). Furthermore, the top CLR species utilized starch and simple carbohydrates as main substrates while EHR species used acetate and RNA-derived inosine and uracil, in addition to amino-acids. The species-level flux simulations indicated that an increased carbohydrate and mucin consumption could lead to higher leucine, isoleucine and proline production (Method). Proline is involved in bacterial and protozoa pathogens and colonisation at the mucosal surface and virulence ^27,28^. Here we find increased levels of indole production in reto-4 in comparison to reto-1 and reto-5, which suggests that enhanced translocation/absorbance across the gut barrier is occurring in reto-4 resulting in depleted levels at the mucosal surface, whereas the opposite would be true of the reto-1 and reto-5 ^29^. Furthermore, production of possibly deleterious gases such as hydrogen sulphide, ammonia and nitric oxide (NO) is dramatically higher in CLR microbiome, suggesting that it might negatively affect health.

As high fibre diet is thought to counter loss of microbiome richness, therefore we simulated the growth rate of the most abundant species in EHR and CLR top abundant species under two fibre-rich dietary regimens (Fig. 2c, Supplementary Table 15). CLR species grows faster due to their high appetite for simple sugars compared to EHR Species. Both fibre-rich diet increased the growth rate of top EHR species, and slightly decreased the growth rate of the top abundant CLR species (Wilcoxon test, p-value < 6×10^−4^). The personalized community-level metabolic modelling based on the top-abundant 25 species in each subject for the CLR and EHR showed the consistency of the flux ratio with the species-level simulations (Fig. 2d, Method). Interestingly, comparison of reto-4 (CLR) with reto-5 (non-westernized high richness reactotype) by applying individual, community modelling and pRSEA, indicated that non-westernized metabolic features are similar to EHR (Fig. 2e, Extended Data Fig. 5b-d). In particular, simple sugar and starch consumption and BCAA production were higher in CLR compared to non-westernized reactotype (reto-5), while BCAAs consumption was lower in CLR.

### Associations of clinical parameters and metabolomics with reactotypes

To evaluate how the reactotypes relate to different microbiome features and to examine a possible association with clinical parameters and the metabolomics, we used a longitudinal Swedish wellness cohort of healthy individuals (86 subjects), followed for a year and sampled 4 times. In view of its accuracy, the random forest method trained based on the healthy samples (n=2567) was used to predict reactotypes in this cohort. Most samples were assigned to reto-1 (247 samples) and reto-3 (84 samples); the other 3 reactotypes were rare (samples <=5; Fig. 3a, Extended Data Fig. 6a, Supplementary Table 16), consistent with global geographic reactotype distribution. Reactotypes had the expected richness (Extended Data Fig. 6b) and enterotype distribution (Fig. 3b), validating conclusions from the previous analyses. A number of functional features were found to be different between reto-1 and reto-3, amenable to the comparison being sufficiently represented in the cohort (Extended Data Fig. 6c). Interestingly, associations with bioclinical parameters were observed: lower richness reto-3 individuals had higher BMI (Wilcoxon test, p-value 0.02, Extended Data Fig. 6d) and waist circumference, higher amounts of low-density lipoprotein (LDL), apolipoprotein B (APOB), fraction of apoB and apoA1 (APOB.A) than higher richness reto-1 individuals (Wilcoxon test, p-value < 0.01, Extended Data Fig. 6e, Supplementary Table 17). This is consistent with the known association of low microbiome richness with physiological state presenting a higher risk of progression towards metabolic syndrome-related pathologies^28^. In addition, a number of metabolomic features were significantly different between reto-1 and reto-3, such as amino acids, amino compounds, xenobiotics and steroids and organic acids (Wilcoxon test, q-value < 0.05, Extended Data Fig. 6f, Supplementary Table 18). Notably, bile acids were higher in reto-1, while carnitine derivatives (glutarylcarnitine, laurylcarnitine, decenoylcarnitine and succinylcarnitine) were higher in reto-3. Additionally, a single individual classified as reto-5 had clinical parameters such as mass of hemoglobin per red blood cell (MCH), average volume of the red blood cells (MCV), markers of liver damage (Gamma Glutaryl Transferase) and kidney function (Creatinine) lowest in the entire cohort (Extended Data Fig. 6g).

### Reactobiome uncovered environment, host and microbiome interactions

As diet is prone to seasonal changes^30^, we examined the reactobiome over a year in the longitudinal Swedish cohort. The reaction richness was similar across the four seasons (Extended Data Fig. 7a), but the abundances of reactions related to energy metabolism varied (Extended Data Fig. 7b, Supplementary Table 19). Reaction abundances of central carbon metabolism were highest in the spring and lowest in the summer (two-way ANOVA test, p-value < 0.001). Similarly, Acyl-CoA oxidase for fatty acid degradation and alpha amylase for starch degradation increased from winter to spring and decreased in summer. An increase in the proportion of calories from carbohydrates in the spring was reported for a US cohort^30^; and it may be that cohort consumed more fats and starch in the spring, leading to an increases in microbes particularly adapted to use them. To gain more insight on seasonal changes of the reactobiome and host physiology, we analysed 1490 plasma metabolite profiles and 41 clinical parameters^20^ for 86 individuals, using multivariate random effects models (MVEM)(Methods). We performed MVEM on each of the plasma metabolite and clinical parameter with 3500 reactions from the reactobiome present in at least 10% of the samples with non-zero variance. 24 associations between six clinical parameters and 21 reactions were found (FDR < 0.05, Fig. 3c, Supplementary Table 20). Seven reactions were significantly associated with C-reactive protein (CRP), a marker for inflammation in the body (FDR < 5×10^−4^) whilst particularly interesting are hydrogen peroxide (H2O2) and NADPH production, related to oxidative stress. Additionally, lipid production was positively linked to the level of cystatin C (FDR = 7×10^−07^). Also, 4-hydroxybenzoyl-CoA thioesterase for benzoate degradation and salicylate 1-monooxygenase for degradation of aromatic compounds in the gut microbiome were negatively associated with troponin T, as an prognostic marker of anthracycline-induced cardiotoxicity^31^. These associations could be related to xenobiotic degradation by the gut bacteria facilitating host detoxification.

We observed 3705 associations between 135 plasma metabolites and 1138 reactions from the reactobiome (q-value < 5×10^−04^, Supplementary Table 21). Amino acids, fatty acids, organic acids and xenobiotics had high connectivity with the reactobiome (Extended Data Fig. 8a, Supplementary Table 22). Reactobiome features connected with amino acids were dipeptidase, kynureninase, nitrilase, tyrosinase, catechol oxidase, homocitrate synthase, isocitrate-homoisocitrate dehydrogenase and glyceraldehyde-3-phosphate dehydrogenase were directly linked with plasma amino acids (Extended Data Fig. 8b). Another group of reactions connected with plasma amino acids were among carbohydrate and xenobiotic metabolism as the potential carbon source^32^, for amino acid biosynthesis (Supplementary Table 23). Reactions that directly consumed carbohydrates (i.e. mon-, di-, oligo- and poly saccharides) from the reactobiome were also connected to plasma metabolites. The reactobiome clarifies the microbe-host interactions based on consumption of different carbohydrates by the gut microbiome and the mediated plasma metabolites such as, amino acids, lipids, steroids and fatty acids (Supplementary Table 24, Fig. 3d).

## Discussion

We developed the MIGRENE Toolbox as a platform to integrate the microbial non-redundant gene catalogues together with metagenome species to generate MAGMA and community-level models to apply them in personalized microbiome studies. The reactobiome as a new representation of the gut microbiome, revealed the novel facets of the landscape of the individual microbial composition and classified the global landscape of the human gut microbiome in healthy individuals to five reactotypes. Moreover, the reactobiome can be applied to overcome the challenge of unclassified metagenome species that could contribute to gut dysbiosis. More than 50% of the MSPs are taxonomically unclassified at the species level but the reactobiome could overcome the contribution of the unknown species in dysbiosis, by providing a functional profile of the species. The new metagenome assembled genomes could even have this issue in a higher degree and the reactobiome could be applied to investigate their functional features.

Within the five reactotypes, three reaction richness groups were identified, introducing a new intermediary group compared to the gene richness classification of microbiome samples based on a single gene count threshold ^3^. The reactotypes integrate the key enterotype groups ^1^, and also gave a more detailed view of the transitions between the three enterotypes ^2^ *via* two additional reactotypes, by assigning important metabolic functional features.

Reto-2 and reto-4 had the highest enrichment in BCAAs and BCFA metabolism, which are both associated with metabolic and inflammatory diseases^33,34^. The Chinese cohorts are dominant in reto-2 and low richness reto-4 (CLR) and these features might be related to the high prevalence of pre-diabetes (35.7%) and diabetes observed in China ^35,36^. Here we would suggest that reactotype 4 is at increased susceptibility to dysbiosis and the associated inflammatory disorders. Interestingly, high richness European individuals (EHR, reto-1) and non-westernized populations (largely reto-5) had common metabolic features. Part of the similarity might be related to diet as suggested by biomass-guided modelling under different nutrient availabilities. Notably, a high fibre-based diet increased the growth rate of high abundance bacteria in the EHR and non-westernised populations, while it decreased that of CLR. The reactobiome analysis indicated that individuals from same region could be classified into different reactotypes, while these individuals had discrete clinical data.

The reactobiome revealed a strong connection between the gut microbiome metabolic activity and compositional changes with host physiology, which could be used for systematic investigation of the environment-microbe-host interactions and adaptation. For example, several studies have shown the relationship between H2O2 and inflammation^37,38^ but here we find that this association could be due, at least in part, to gut bacterial metabolic activity. Moreover, the reactobiome revealed the comprehensive relationship between the plasma metabolites and microbiome metabolic activity such as, degradation of carbohydrates by the gut bacteria that could impact on metabolite levels in the host. As the human host can adapt itself to seasonal and dietary changes^39,40^, here using the longitudinal cohort, we explored the metabolic changes in the gut microbiome in response to changes in environmental perturbations through the reactobiome, which could lead to gut microbiome adaptation and consequently affect host physiology. Therefore, this study starts to decipher the complex interactions between host and microbes, which may help identify future therapeutic strategies targeting the gut microbiota.

## Methods

### Metagenomics species pan-genome (MSP) and completeness estimation

The updated Integrated Gene Catalogue of the human gut microbiome (IGC2)^21^ was used to generate the generic metabolic model. The updated MSP information was used here to reconstruct the species-specific models ^18^. All the metagenome samples have been previously mapped to the IGC2 and profiled for the MSP abundances. MSP completeness was estimated using two conserved single-copy marker genes (SCM) sets that have been previously determined for all bacterial and archaeal finished genomes^41^. The sets consist of 139 (bacteria) and 162 (archaea) SCM that were found to occur only once in at least 90% of all genomes. Species completeness score was estimated as the proportion of SCM genes found in the MSP using hmmsearch and HMMER^42^ formatted databases for archaeal and bacterial SCM. In order to compare the completeness of the generated MAGMA vs nonfunctional MSP models, we used two-sided Wilcoxon signed rank test.

### MIGRENE toolbox; Integration of bacterial gene catalogue into metabolic model and microbiome reference GEM generation

MIGRENE toolbox was utilized for the integration of IGC2 into metabolic model. First, the bacterial gene catalogue was restructured and checked using *checkCatalog*.*m* function. *convertCatalogAnnotation*.*m* function converted the KO annotation from catalogue into KEGG reaction ID. If the mapping ID file is not assigned as an input argument, it automatically downloads the information for ID conversion from KEGG API and saves in directory “data” where the MIGRENE toolbox is located. Second, a generic annotated metabolic model was provided based on ModelSEED ^43^and KBase^44^ to maximise the coverage of catalogue KEGG gene annotation. We corrected the directionality of the reactions, defined 165 exchange reactions and their corresponding transport reactions, biomass (Supplementary Table 25), compartmentalised the model, and removed dead end reactions and metabolites. We also balanced reactions in regard to the molecular formula of the metabolites in the generic model. The metabolites on both sides of reactions were checked for carbon, oxygen, phosphor, sulphur and nitrogen. The generic metabolic model and the IGC2 were used as inputs to the *microbiomeGEMgeneration*.*m* function to generate a gut reference GEM. The Gene-protein-reaction (GPR) association was assigned by integrating the catalogue genes into metabolic model using the KEGG reaction IDs. In case of the annotation file for GEM is not provided, the tool automatically finds the annotations i.e. KO, reaction ID or EC in the model and convert into reaction ID based on the latest version of KEGG IDs. The function adds fields “grRules”, “genes”, “rules”, “geneNames” and “rxnGeneMat” into the GEM to be compatible with both COBRA^45^ and RAVEN toolbox^46^ structure, so the output models can be used for any functions provided in both toolbox.

### Diet allocation for GEMs

Diet allocation for GEMs was built using the USDA food composition database (https://ndb.nal.usda.gov/ndb/). Based on the molecular weight for each micronutrient, the mmol/gDW was calculated and divided per 24 hours (mmol/gDW*hour). Food items that had a missing value for an amino acid but contained protein were replaced by the average of the amino acid for the corresponding food group. This value was then divided by the total average of all foods in the database. Four different diets (high-protein-plant based, high-protein omnivorous, high fibre-plant based and high fibre omnivorous) were formulated based on macronutrient composition^47,48^. Each diet is an average diet based on a three-day meal plan in which each day consisted of three main meals and a snack. All ingredients for each meal from the four different diets were similar, and certain ingredients were swapped for alternatives to fit the specific diet. The composition of each diet is given in appropriate portion sizes and normalized based on 2000kcal a day. Each diet includes information on the macronutrient content such as the energy consumption, carbohydrates, protein, fats and fibres. In addition, each diet contains information on the micronutrient content such as vitamins, minerals, amino acids and ions. The diets were converted to mmol/gDW*hour using *USDAcreatingDiet*.*m* function and added to MIGRENE toolbox.

### MIGRENE toolbox; calculation of reaction score and species-specific model (here MAGMA) generation

Before generating species-specific models, MAGMA (<u>M</u>etagenome species <u>A</u>ssembled <u>G</u>enome-scale <u>M</u>et<u>A</u>bolic models), the gut reference GEM was constrained based on high fiber animal diet using *DietConstrain*.*m* (Option diet number 2). Acetate and Lactate were added as popular carbon sources for bacteria for Constraint-based modelling. This approach was applied to quantitate the reaction rate and calculate reaction fluxes using COBRA Toolbox^45^. Biomass as the main objective function was provided as sets of metabolic tasks to check the functionality of models. Additionally, phenotype of species were manually checked and annotated based on JGI-GOLD phenotype data (organism metadata) (https://gold.jgi.doe.gov/organisms). The bibliomic data of phenotypic features were provided in a data structure with four categories: “bacteria” a cell array listing the name of the bacteria, “rxn” that list the name of the reactions having bibliome data, “value” a matrix including 0 (no information), 1 (consumed), 2 (produced), -1 (not-consumed) and -2 (not-produced) by the corresponding bacteria and “aerobeInfo” a cell array provides oxygen requirement for growth of bacteria (aerobe, anaerobe or facultative).

We created a reaction profile for each species based on mapping absent/present genes in MSP using the gut reference GEM (*MetagenomeToReactions*.*m* function in the toolbox). The function also filters the gene vector and gene rules in reference model for each species as a first step for MAGMA generation. Following, the reaction states for all the MSPs and taxonomy information were collected (*GenerateMSPInformation*.*m)*. Reaction scores for each MSP (bacterial species) were calculated based on the bacterium gene profile and the frequency of each reaction in different taxonomy levels of the corresponding bacterium (*MetaGenomicsReactionScore*.*m)*. The function also calculates a threshold (the lowest nonzero frequency) for each taxonomy level for filtering the reactions while generating the context specific GEMs. *contextSpecificModelGenertion*.*m* and then *contextSpecificModelTune*.*m* were used to generate the MAGMA models. For the gap filling, we pruned the constrained reference GEM from the lowest reaction score and minded the gaps to keep the model functional based on biomass objective function and the threshold. Therefore, the gap filling from the reactions starts from the highest frequency in the closest taxonomy level of the species. The latter function tuned the models by pruning the exchange and transport reactions, dead end metabolites, adding compartments, model description and correcting KEGG metabolite ID and metabolite formula. It also calculated the important information for gap filling percentage at different levels: i.e. taxonomy proximity based, further taxonomy level, not annotated (without gene-protein-reaction relation) and the total gap filling percentage. The growth rate for each model were tested for the diets using constraint-based modelling. Additionally, the structure, connectivity (*GEMtoBiPartiteGraph*.*m*) and carbon balance of the functional models were investigated against other publicly available GEMs resources ^10,22^.

### MIGRENE toolbox; generating personalized metabolic microbiome model

A reaction pool based on all the reactions in MAGMA models and the MSP abundance of the corresponded MAGMAs were used for personalized metabolic microbiome model. Reaction richness profile that provides gut microbiome reaction composition for individuals was calculated using *RxnRichnessGenerator*.*m* function. The abundance of each MSP was considered for the reactions of the MSP in the sample. So, we generated a matrix (*n*×*m*; *n*, number of reaction pool and *m*, number of MSPs) for each sample. each matrix was converted to a binary matrix and then to a binary vector that shows absent/present of each reaction in the samples; present means the reaction was captured at least in one MSP in the sample. The count of the present reactions provided reaction richness of the sample. Based on the sum of the abundance for the reactions, relative reaction abundance for the samples was calculated *(ReactionAbundanceGenerator*.*m)*. The reaction abundance profile represents the summation of the respective reaction abundance in all MSPs divided by summation of all reaction abundance in the sample. Then the Reactobiome was determined which is the number of the reactions in the gut and normalized per 500 bacteria for each individual, named count per 500 bacteria (CPF) (*CPFGenerator*.*m)*.

The personalized reaction set enrichment (pRSE) is overrepresentation of each pathway in sample *(pRSEGenerator*.*m)*. the enrichment analysis was performed for 153 KEGG metabolic pathways (Supplementary Table 26). We calculated the prevalence of the enriched metabolic pathways (p-value <0.05) in each sample (*pRSEGenerator*.*m)*. The overrepresentation of the pathways was calculated based on Hypergeometric test and the binary vector. Community models for 2911 samples were generated *(MakeCommunity*.*m)*. Top 25 MAGMAs per samples were selected for community modelling due to computational power required to reconstruct and simulate community models for 2911 samples. The name of MSPs as prefix were added into the reaction names in the GEMs before generating a combined S matrix to keep the species-specific GEMs separated in the community models. Additionally, two compartments named *[lu]* for intestinal lumen and *[fe]* for secreted bacterial and remaining food-derived metabolites were added to the community models. *[lu]* includes represent the gut microbiome and includes all the bacterial GEMs. All the bacterial exchange metabolites were connected to *lu* by designing an extra exchange reaction layer between the *lu* and the bacteria. *FoEx* irreversible exchange reactions transfer food-derived metabolites to *lu* and distributed in the whole community. *FeEx* irreversible exchange reactions transfer the metabolites (bacteria/food drived) into *[fe]* in order to leave the community toward blood or faeces. A community biomass objective function was designed. It is composed of all the bacterial biomass in the community where the abundance of bacteria was used as stoichiometric coefficients. We performed flux balance analysis using COBRA toolbox for the communities where community biomass objective function was maximized.

### Clustering of reactobiome profile, modelling and features of Reactotypes

Dirichlet multinomial mixtures (DMM) method^23^ was performed using “DirichletMultinomial” R package for clustering of reactobiome profile. The number of clusters (reactotypes) were determined by fitted Dirichlet mixtures and Laplace approximation to the model using *dmm* and *laplace* functions. The robustness of the reactotypes were investigated using supervised machine learning models and cross validation using “caret”, “e1071” and “randomForest” R packages. The parameters (50 folds repeat 10 times) for *train* function were set using *rfeControl* function. The collation of resampling results was performed by *resamples* function. We subsequently used wellness cohort as a test set and trained random forest based prediction model for reactobiome profile. We used random forest model to estimate variable importance in reactotypes using *varImp* function in caret package. *pr*.*curve* and *roc* functions computed the area under the precision-recall curve and built receiver operating characteristic (ROC) curve, respectively.

We metabolically investigated the differences among the reactotypes using reactobiome and reaction abundance. non-parametric Kruskal-Wallis test was used for reactobiome profile as negative Binomial distribution and the test was followed by Dunn’s multiple comparisons test for pairwise comparison of reactotypes. Among five reactotypes, we estimated cohen’s d effect sizes for on-sided Wilcoxon signed rank tests of reactobiome in two different reactotypes. Based on estimated effect sizes (Cohen’s d >= 0.5), the significantly enriched reactions were selected and defined as “reactotypes-enriched” reactions. Cohen’s d < 0.5 were selected as depleted reactions. Besides, we employed Multidimensional scaling (MDS) to identify how samples of reactotypes were clustered based on reactobiome profile. we also calculated centroids and standard deviations of samples of the sixteen countries and projected to the MDS plot.

### Enterotype assessment

Abundance profiles of MSPs over total 2,911 samples were summed up into abundances of total 316 genera. Then this genus-level profile was clustered into three representative group by unsupervised clustering method, Dirichlet multinomial model^23^ i.e. R DirichletMultinomial package. Based on enriched genera of each representative group, we analysed the characteristics of microbial compositions and named by most enriched genera of respective groups, such as Bacteroides, Prevotella, and genera belonging to Firmicutes. After identification of enterotypes enriched in Bacteroides (ET-Bacteroides), Prevotella (ET-Prevotella), and Firmicutes (ET-Firmicutes), fraction of three enterotypes were investigated over 2,911 samples.

### Microbiome-host and seasonal changes association studies

In order to investigate the association for each reaction, we filtered the reactions <0.01% of reactobiome composition and prevalence > 10%. The multiple covariates between reactobiome profile and host clinical and plasma metabolites in the population-scale longitudinal study were subjected to multivariate regression, considering random effects of individuals by linear mixed-effect models (multivariate random effects models) using MaAsLin2 R package (https://huttenhower.sph.harvard.edu/maaslin). Significant associations (adjusted p-value <0.01) were visualized using circlize R packages^49^ for circus plot and Cytoscape 3.7.2 ^50^ for network representation. The abundance of the reactions was analysed using two-way ANOVA to detect seasonal changes in individuals (n=86) in a combination of two independent variables i.e. data points and subjects.

## Data availability

The 1333 reconstructed MAGMAs are available in www.microbiomeatlas.com.

## Code availability

The MIGRENE toolbox used to reconstruct MAGMA, quality control, generating reactobiome, reaction abundance, calculating reaction richness, personalized and community modelling, pRSE and diet allocation, with several tutorials and testing datasets, can be found at our GitHub repository https://github.com/sysbiomelab/MIGRENE. The toolbox can be applied to any microbial gene catalogues and metagenome species for GEMs reconstruction. In addition, any public microbial GEMs, together with the abundance, can be utilized for reactobiome, reaction richness and abundance, and community modeling.

## Acknowledgements

This study was supported by Engineering and Physical Sciences Research Council (EPSRC), EP/S001301/1, Biotechnology Biological Sciences Research Council (BBSRC) BB/S016899/1, Science for Life Laboratory, the Knut and Alice Wallenberg Foundation, the Erling Persson Foundation and King’s College London. We thank the Swedish National Infrastructure for Computing at SNIC through Uppsala Multidisciplinary Center for Advanced Computational Science (UPPMAX) under Project SNIC 2019/3-226, SNIC 2020/6-153, SNIC 2020/5-222.

## Author contributions

SAS, SDE and GB conceived the project. GB developed MIGRENE toolbox, generated MAGMA and community models, reactobiome and reactotypes and host-microbiome interaction. SL performed enterotype analysis. MA, ELC, FPO and NP provided MSPs information. BZ assisted in metabolic model curations and diet allocation algorithm. GB, SDE and SAS wrote and drafted the manuscript. SL, LAE, DLS, MA, SDE, LN, JN, GP and MU provided critical feedback on the data and manuscript. All authors read, edited and reviewed the manuscript.

## Extended Data figure legends

**Extended Data Fig. 1.**
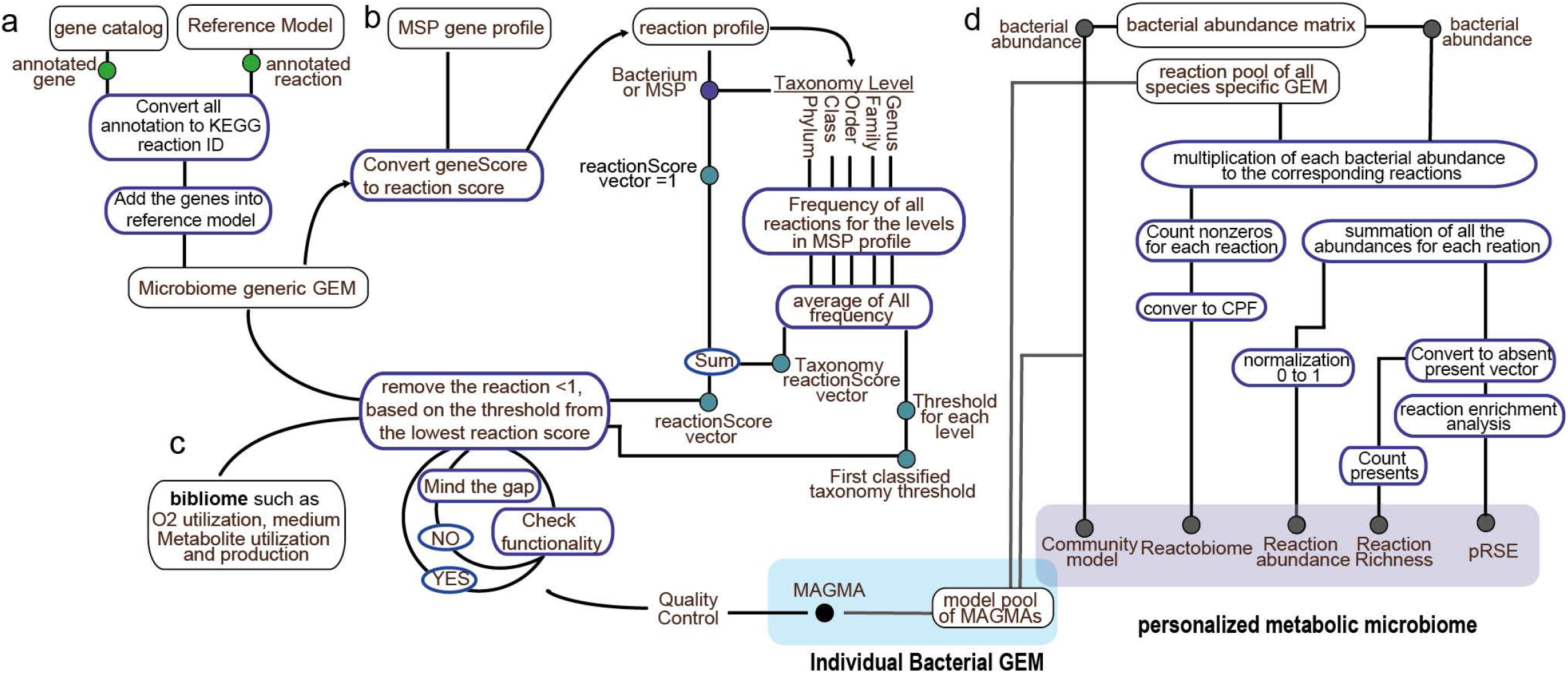
The MIGRENE Toolbox. **a**, the software allows for integration of different microbial gene catalogues with any available functional annotations to construct the specific microbial metabolic reference network. The initial step in the workflow is generating a microbiome generic GEM, using integration of the non-redundant microbial gene catalogue and a reference model such as KBase or ModelSEED ^44,52^. **b**, the reaction profile for the metagenome species, in this case MSPs, will be generated based on mapping to the reference model. In addition, reaction frequency and threshold for the MSP based on the taxonomy level will be calculated. The reaction profile and frequency together will construct the reaction score for the MSP. Additionally, a threshold for each reaction frequency is determined. **c**, based on the updated reaction frequency vector, bibliomic data, and threshold the species-specific GEM based on the MSP-level information is reconstructed. The process is run iteratively by benchmarking the functionality of the model for growth. The generated GEM based on the MSP-level information, are called <u>M</u>SP <u>A</u>ssociated <u>G</u>enome scale <u>M</u>et<u>A</u>bolic models (MAGMA). **d**, the analysis part of the toolbox uses the MAGMA models, but any other microbial GEMs could be integrated into the metagenomics data for “personalized metabolic microbiome analysis”. This generates four outputs at a personalized level; the reactobiome and reaction abundance, reaction richness, community modelling, and personalized reaction set-enrichment (pRSE) (see Methods). Reaction abundance is computed as sum of the respective reactions from corresponding gene abundance; the reactobiome is the number of the reactions normalized per 500 bacteria for each individual, named count per five hundred bacteria (CPF) (Method). CPF removes the high heterogeneity of bacterial presence in each individual (3 to 385 MSPs in the HGMA samples) and technical biases of sequencing ^53^. It facilitates the comparison of reaction counts between the samples from different cohorts. Reaction richness determines the reaction composition in each metagenome sample. Community modelling performs constraint-based analysis on MAGMAs, the metabolic interactions and productions from the microbial community. pRSE captures the over- and under-representation and prevalence of metabolic pathways.

**Extended Data Fig. 2.**
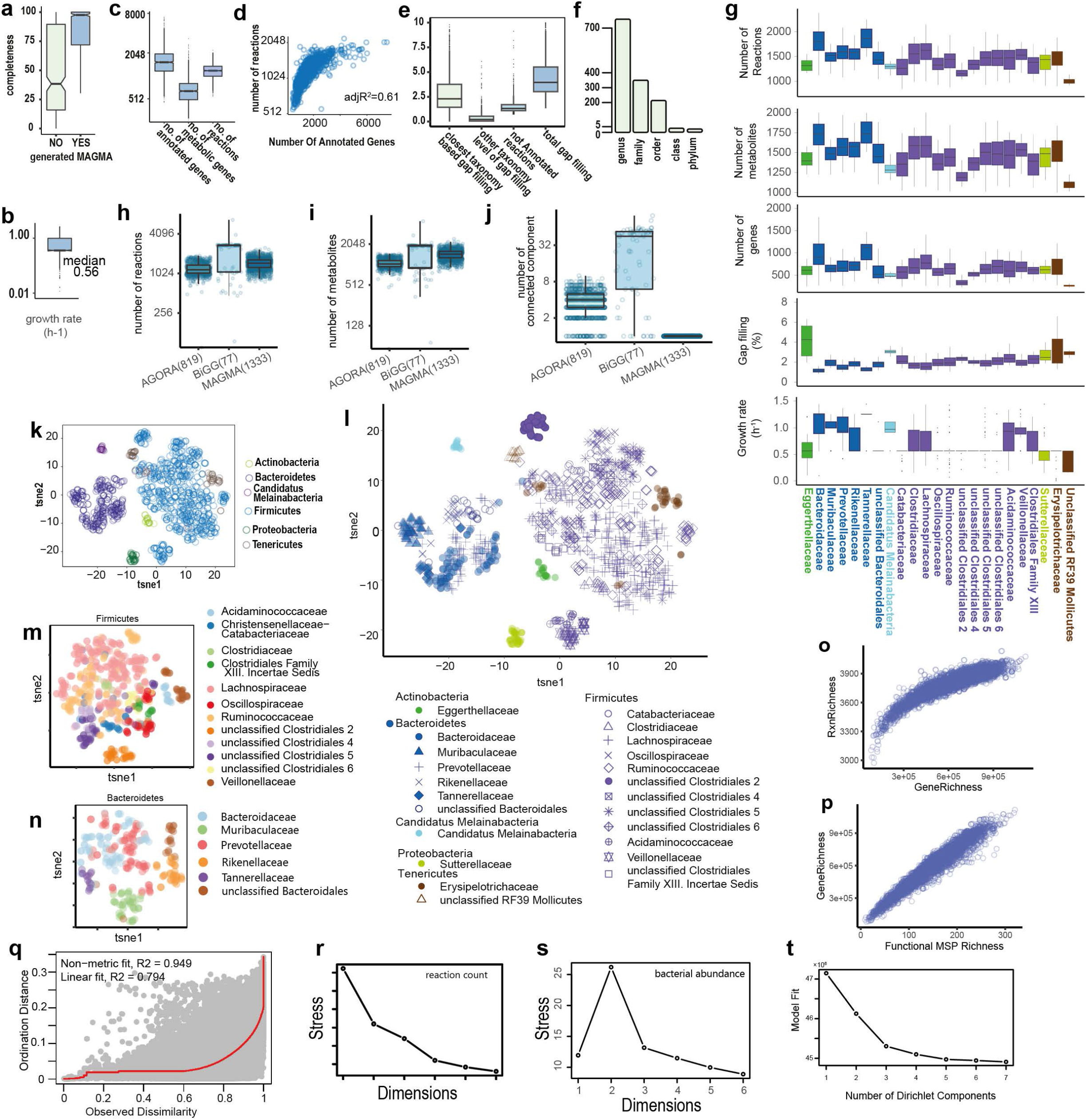
MAGMA features and global landscape of the reactobiome and reaction richness in healthy samples. **a**, boxplot for completeness of MSPs yielding functional and non-functional MAGMAs. **b**, the simulated growth rate (hr^-1^) of MAGMAs constrained by western diet in anaerobic condition. **c**, Boxplot for metabolic and annotated genes and number of reactions for MSPs with functional MAGMAs. **d**, Scatter plot for number of reactions in MAGMAs and the number of genes in MSPs. **e**, Boxplot of the gap filling percentage at different levels: i.e. taxonomy proximity based, further taxonomy level, not annotated (without gene-protein-reaction relation) and the total gap filling percentage. **f**, Bar plot shows the number of MAGMAs per taxonomy level of gap filling. **g**, MAGMA features for the families with more than 10 MAGMA members. A negative correlation between percentage of gap filling and the number of annotated genes in MSPs was observed. **h**, reactions, **i**, metabolites and **j**, connected components in GEMS in other sources (AGORA, BiGG) and MAGMA; number of the GEMs used for comparison is indicated in parentheses. **k**, Visualization of MAGMA clusters with t-SNE at phylum level (only the phyla with more than 10 MAGMA members were used). **l**, visualization of MAGMA clusters with t-SNE at family levels (only the families with at least 10 MAGMA were used). Visualization of MAGMA clusters of **m**, Firmicutes and **n**, Bacteroidetes by t-SNE. **o**, Scatter plot of gene richness and reaction richness in the healthy samples (n=2911). **p**, scatter plot of the functional MSP (MAGMA) richness and gene richness in the healthy samples (n=2911). **q**, NMDS Shepard plot of the relationship between the data dissimilarities (x-axis) and the ordination distances (y-axis) by MSP abundance profile. The fit between dissimilarity and ordination distances is indicated by red line. **r**, scree plots for reactobiome profile showing the decrease of ordination stress in dimensions of 1 to 6. **s**, Scree plots for bacterial abundance profile showing the decrease of ordination stress in dimensions of 1 to 6. **t**, increasing number of Dirichlet mixture components (1 to 7) regarding the Laplace approximation

**Extended Data Fig. 3.**
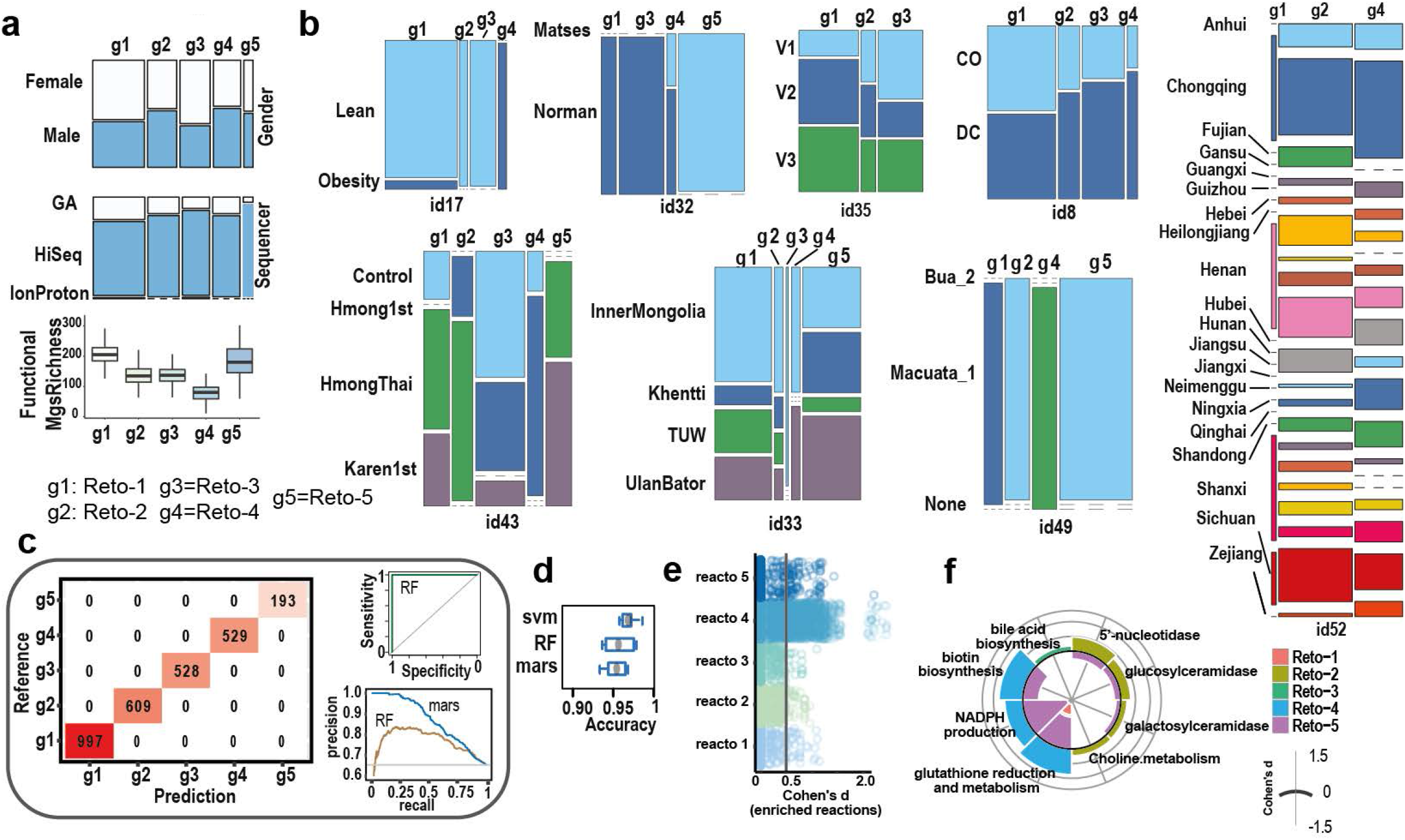
Characterization of reactotypes. **a**, MAGMA richness and gender distribution in reto-1 (g1) to reto-5 (g5). **b**, the distribution of samples in reactotypes 1-5 based on the available metadata in their cohorts. **c**, the predicted number of all healthy samples in each reactotypes were compared to the reference reactotype (obtained from DMM) using the random forest method. AUC=1 in receiver operating characteristic (ROC) curve. Precision-recall curves for multivariate adaptive regression splines (MARS) and random forest (RF) classification methods. **d**, accuracy of the machine learning methods support vector machine (SVM), RF and MARS by resampling. **e**, Cohen’s d effect size of one-sided Wilcoxon tests for reactotype-enriched reactions. **f**, radar plots showing the fraction of functional terms in reto-1 to reto-5, tested by enriched/depleted reactions of the reactobiome based on Cohens’ d effect sizes of Wilcoxon one-sided tests (positive and negative Cohens’ d respectively indicate enriched and depleted reactions).

**Extended Data Fig. 4.**
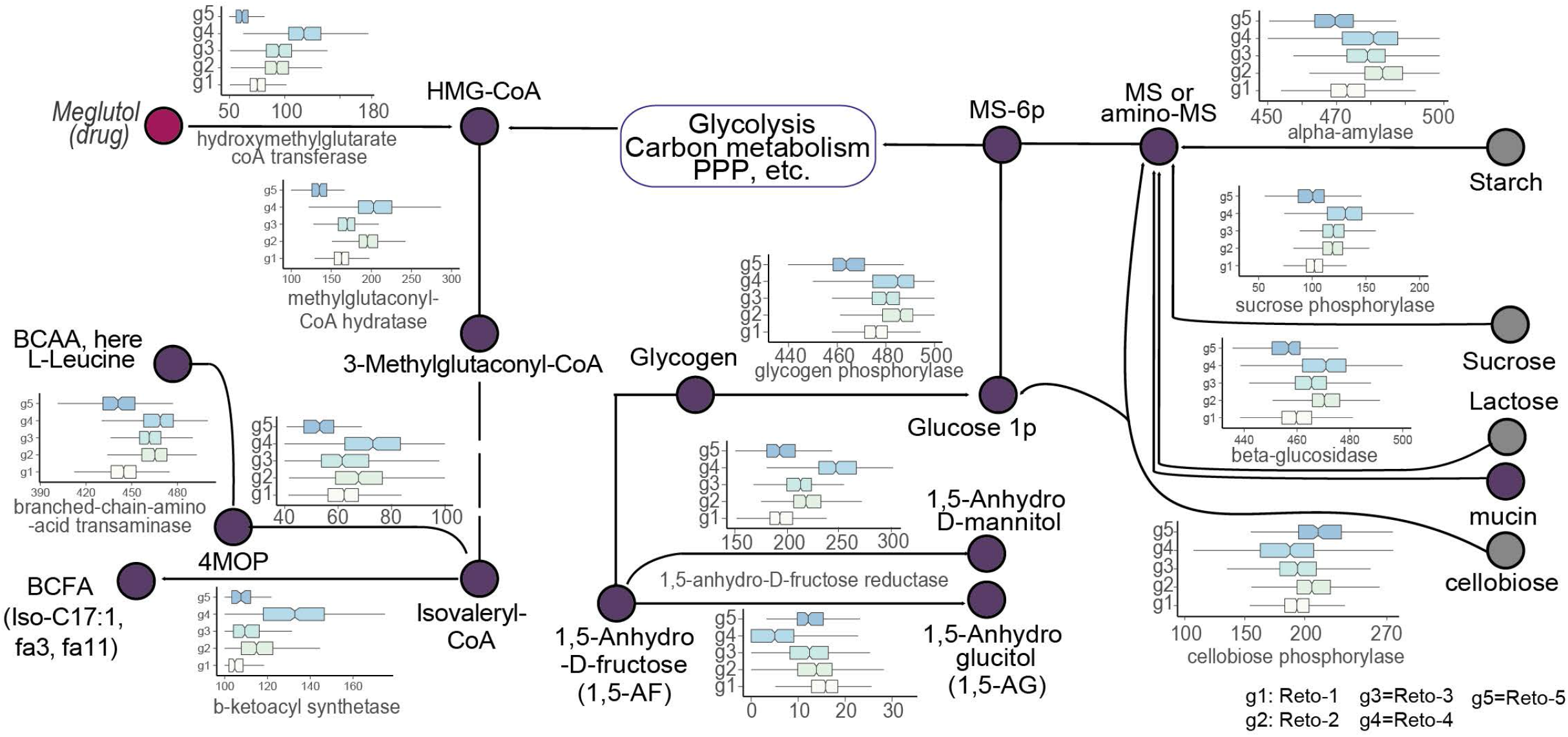
Metabolic features in reactotypes. Connection of metabolic pathways of macronutrients (grey circles) and antilipemic drug (Meglutol) as features in reactotypes. g1 to g5 indicates reto-1 to reto-5, respectively.

**Extended Data Fig. 5.**
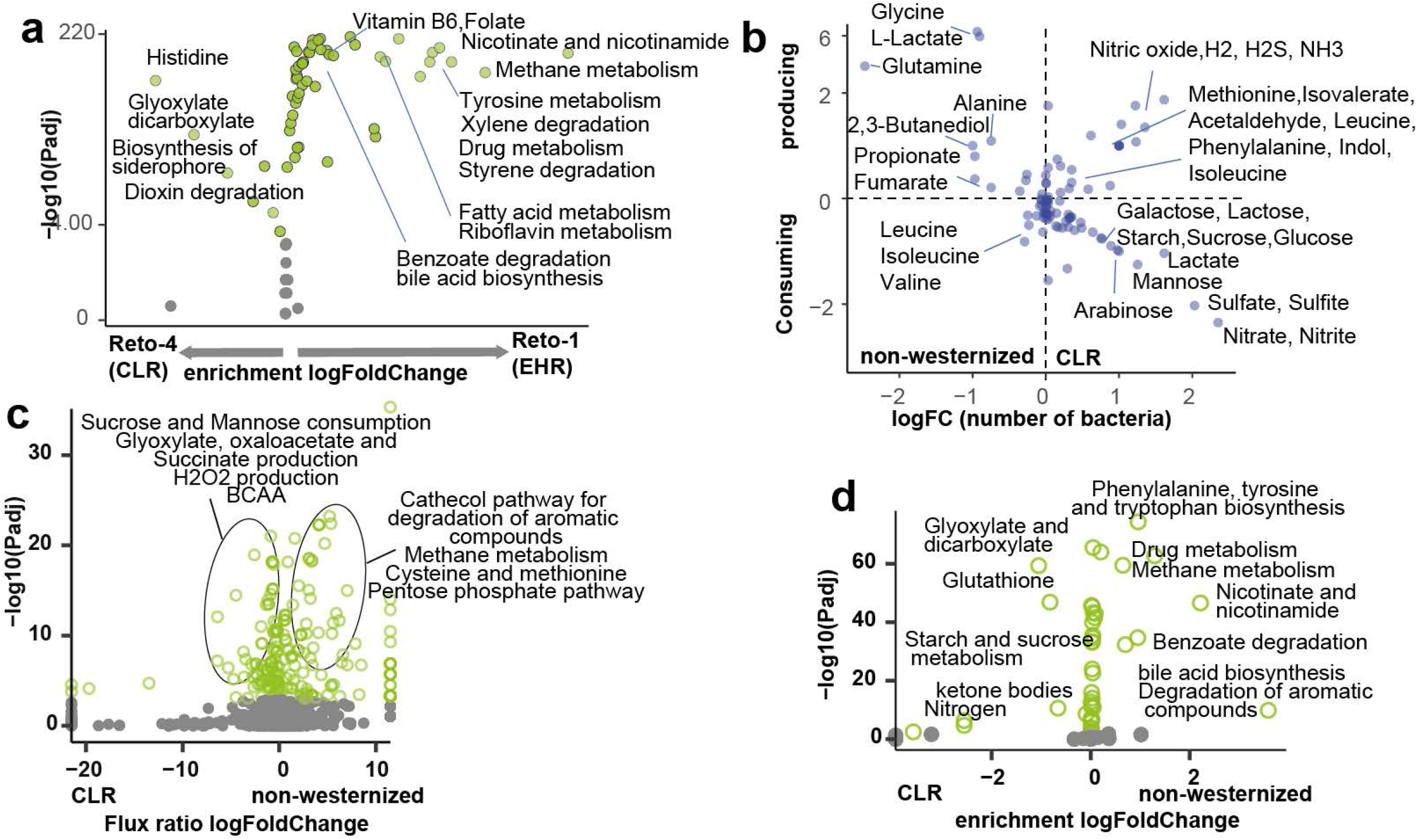
Diferences of gut microbiome metabolism between Chinese low richness (CLR, ret-4), and high richness (EHR, ret-1) reactotypes. **a**, differences of personalized enriched pathways between EHR (reto-1) and CLR (reto-4) groups (FDR < 0.001, Wilcoxon test. green dot: substantially different). pRSEA was performed to calculate overrepresentation of metabolic pathways in each sample. **b**, the production/consumption log fold change of non-westernized vs CLR (Y axis) and the log fold change of number of top 50-ranked abundant bacteria in each group producing/consuming the corresponding metabolites. One of the main common features between non-industrialized and CLR population is the sugar appetite in the high abundant bacteria. BCAAs isoleucine and leucin and indole and isovalerate are produced in CLR more than non-industrialized gut microbiota while production of propionate and lactate are higher in the non-industrialized group. Production of gases such as hydrogen, hydrogen sulphide and NH3 are dramatically higher in the top industrialized bacteria compared to non-industrialized population. additionally, the signaling molecule nitric oxide (NO) is also released by industrialized gut microbiome more than non-industrialized regions. **c**, the differences of flux ratio between personalized gut bacterial community of non-westernized vs CLR (FDR < 0.001, Wilcoxon test. green dots: substantially different). **d**, differences of personalized enriched pathways between two populations (FDR 0.001, Wilcoxon test. green dots: substantially different. pRSEA was performed to calculate overrepresentation of metabolic pathways pathway in each sample.

**Extended Data Fig. 6.**
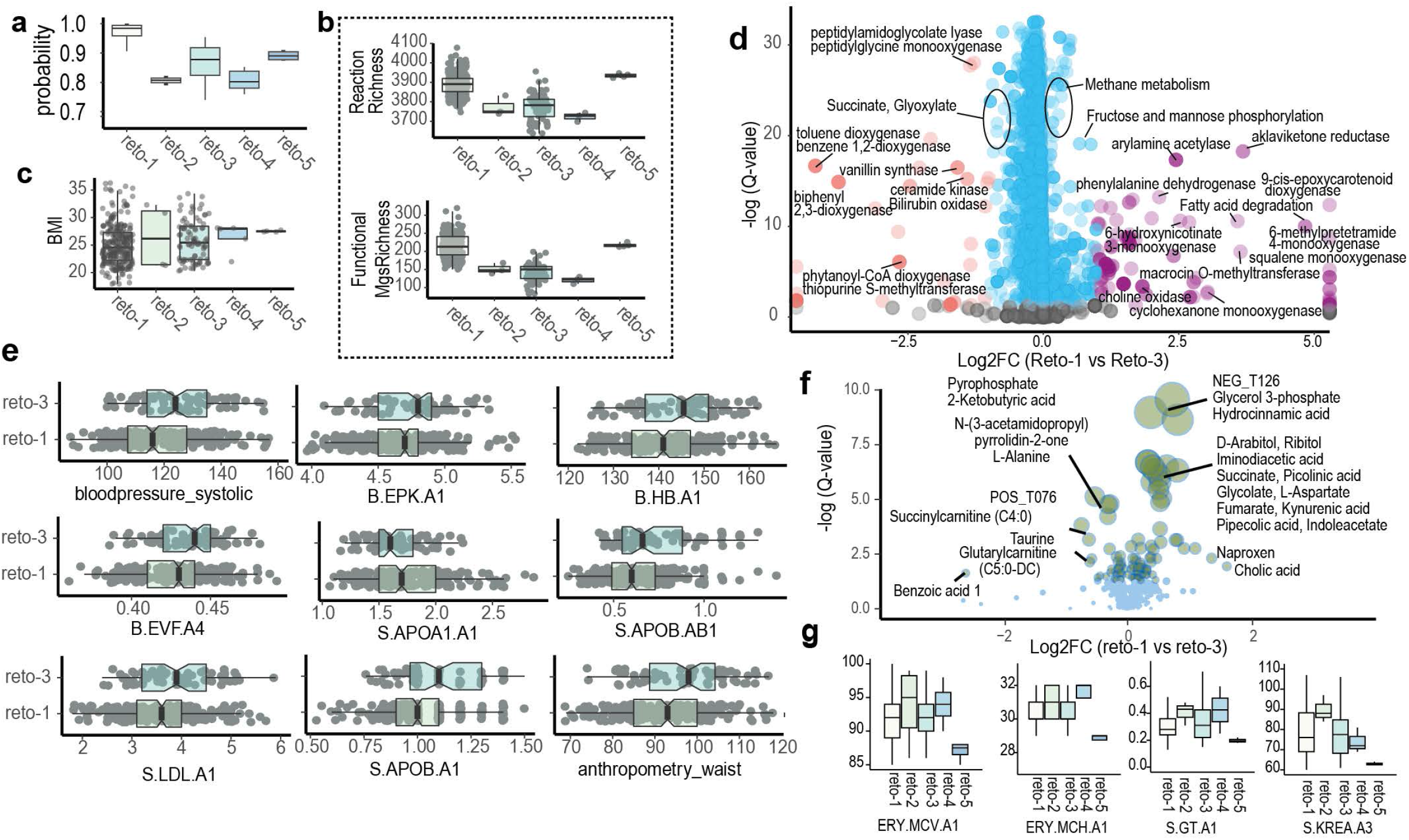
Reactotypes in Swedish wellness cohort. **a**, the random forest predicted probability for reactotypes. **b**, the boxplot of reaction richness and MSP richness (MSPs with functional reconstructed model). **c**, BMI in wellness cohort reactotypes 1-5. **d**, reactobiome differences between reto-1 (n=247) vs reto-3 (n=84) at FDR 0.01, Wilcoxon test with Benjamini-Hochberg correction. Red dots: highly reto-3-enriched reactions (Log2 FC < - 1); purple dots: highly reto-1-enriched reactions (Log2 FC > 1); blue dots: significantly enriched; grey dots: not significant **e**, boxplot of clinical parameters in reto-1 and reto-3. Erythrocyte volume fraction (B.EVF.A4), Erythrocyte particles (B.EPK.A1) and Hemoglobin (B.HB.A1) related to erythrocytes and blood pressure (bloodpressure_systolic) were higher in reto-3 with lower richness. **f**, changes of plasma metabolites between reto-1 vs reto-3 in wellness cohort green circles: significantly different (Wilcoxon test, FDR < 0.05). **g**, boxplot of clinical parameters in wellness cohort for reto-1 to 5. S.KREA.A3: Creatine; S.GT.A1: Gamma-glutamyltransferase; ERY.MCV.A1: Mean corpuscular volume (MCV); ERY.MCH.A1: mean corpuscular hemoglobin.

**Extended Data Fig. 7.**
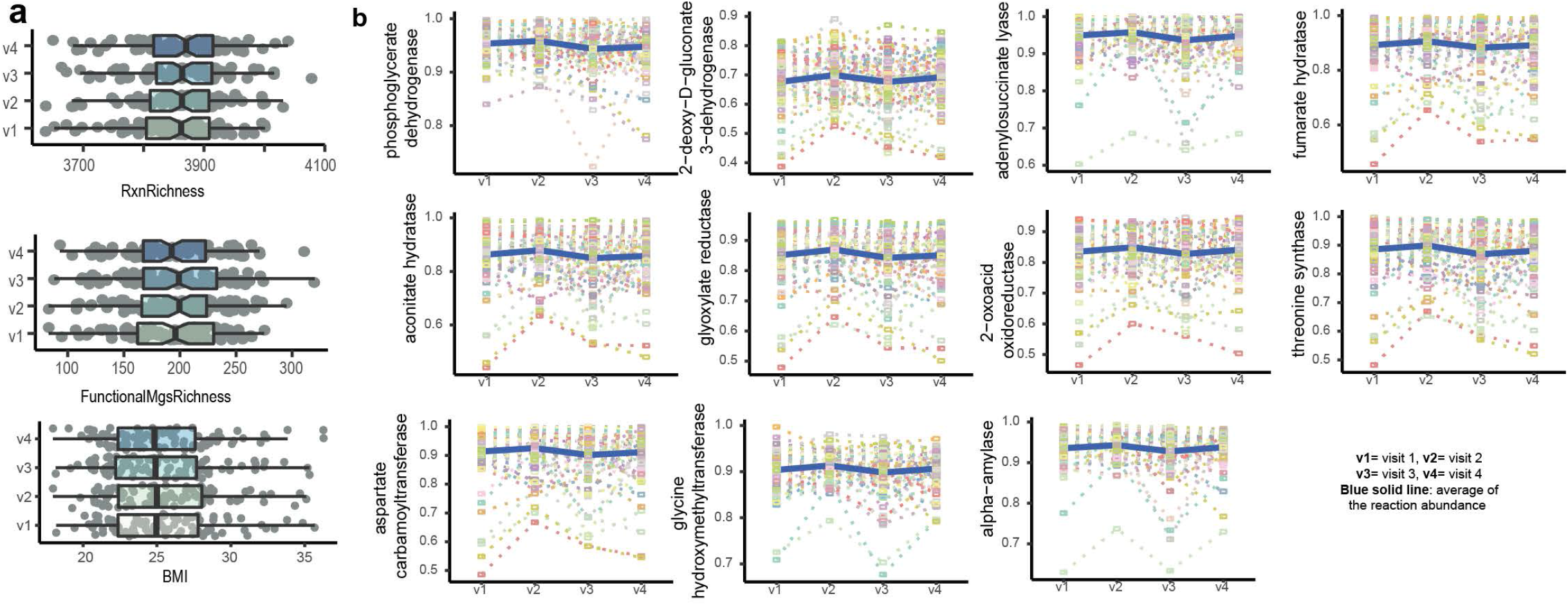
Reactobiome and richness variation across the four visits (v1-4). **a**, boxplot of different richness profiles in wellness cohort in four visits. **b**, reactobiome changes across the four visits for 86 individuals using two-way ANOVA (p-value < 0.01). Each individual in time points were shown with different box colours and the dotted line to connect the individual across the time. v1: winter; v2: spring; v3: summer; v4: autumn.

**Extended Data Fig. 8.**
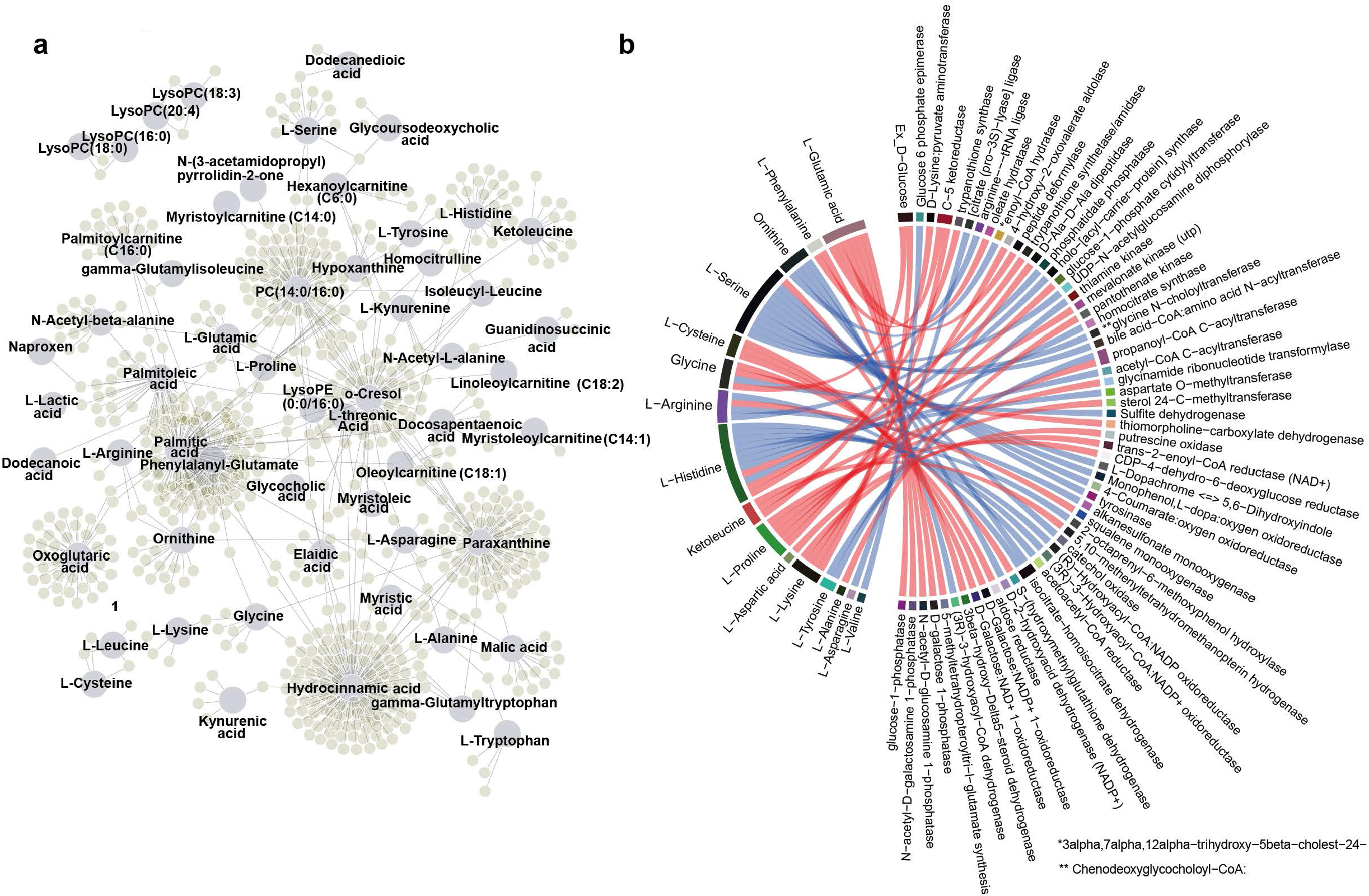
Interplay of host plasma metabolites with variation of the gut reactobiome. **a**, the network indicates the association between variation of plasma metabolites with the reactions in the gut reactobiome at FDR < 0.01, using multivariate random effects model. **b**, chord diagram shows the reactobiome reactions that are linked to plasma amino acids at FDR < 0.01, using multivariate random effects model.

## Supplemental figure legends

**Supplementary Fig. 1**| **Galactose metabolism**. Reactions enriched (shown in red) in CLR relative to EHR (compared with high richness) at FDR 0.01, Wilcoxon test.

**Supplementary Fig. 2**| **Amino sugar and nucleotide sugar metabolism**. Reactions enriched (shown in red) in CLR relative to EHR (compared with high richness) at FDR 0.01, Wilcoxon test.

**Supplemental table legends**

**Supplementary Table 1**| Description of 1333 MAGMA including taxonomy, number of reactions, number of metabolites, number of genes, number of reactions, the level of gap filling, percentage of gap filling, biomass flux.

**Supplementary Table 2**| comparing the topology of MAGMA models with AGORA and BiGG GEMs. The number of connected Components, metabolites, reactions and carbon balanced were compared.

**Supplementary Table 3**| Description of the cohorts from Human Gut Microbiome Atlas (HGMA) datasets used in this study including the id, total number of samples, number of the healthy samples, sequencing platform and geography.

**Supplementary Table 4**| 2911 healthy samples including cohort id, MSP richness, gene richness, reaction richness and functional MSP (MAGMA) richness.

**Supplementary Table 5**| reactobiome profile of 2911 healthy samples for 4397 reactions

**Supplementary Table 6**| MSP abundance profile of 2911 healthy samples.

**Supplementary Table 7**| the members of five reactotypes (reto-1 to reto-5) contains sample id, dataset id, gender, geography, enterotype, age and reactotype.

**Supplementary Table 8**| reaction comparison among reactotypes by non-parametric tests Kruskal-wallist test and Dunn’s multiple comparisons and random forest for feature selection for 3516 reactions annotated by EC, KEGG reaction ID and KO.

**Supplementary Table 9**| reactotype-enrichement/depletion of 3516 reactions were tested by Cohen’s d effect size of one-sided Wilcoxon tests.

**Supplementary Table 10**| reaction abundance profile of 2911 healthy samples for 4397 reactions.

**Supplementary Table 11**| comparing reaction abundance involved in sugar and amino sugar metabolism and mucin degradation in EHR (Reto-1) vs CLR (Reto4) using Wilcoxon two-sided tests adjusted p-value and FDR.

**Supplementary Table 12**| comparing reactions abundance in Reto-5 vs CLR (Reto4) using Wilcoxon two-sided tests adjusted p-value and FDR.

**Supplementary Table 13**| top 50-ranked MSPs significantly abundant in reto-1 and reto-4.

**Supplementary Table 14**| simulation of growth rate (h-1) of top 50-ranked bacteria of reto-1 and 4 based on the maximization of the biomass in the presence of high protein and fibre omnivorous and plant based diets included Lactate and Acetate.

**Supplementary Table 15**| simulation of top ranked bacteria constrianed by high fibre plant based, high fibre omnivorous and high protein omnivorous diets.

**Supplementary Table 16**| reactotype analysis on 344 samples from Swedish cohort including predicted reactotype, probability of predicted reactotype, Enterotype group, gene and reaction richness, MSP and MAGMA richness, BMI and age.

**Supplementary Table 17**| clinical data significantly enriched in Reto-1 or Reto3 (Wilcoxon two-sided tests adjusted p-value <0.01) of Swedish cohort.

**Supplementary Table 18**| metabolomic data significantly enriched in Reto-1 or Reto2 (Wilcoxon two-sided tests adjusted p-value <0.01) of Swedish cohort.

**Supplementary Table 19**| reactions significantly different among four seasons by ANOVA test and multiple comparisons (p-value < 0.01).

**Supplementary Table 20**| Statistics of associations between reactobiome and clinical parameters. We identified 24 significantly association by multivariate random effects model. It shows corresponding coefficient, p-value, and FDR of significantly associated clinical parameters.

**Supplementary Table 21**| Statistics of associations between reactobiome and plasma metabolites. 3705 associations between 135 plasma metabolites and 1138 reactions from reactobiome were observed by multivariate random effects model. it shows corresponding coefficient, p-value, and FDR of significantly associated plasma metabolites.

**Supplementary Table 22**| number of reactions in reactobioms associations with plasma metabolites.

**Supplementary Table 23**| reactobiome connected with plasma amino acids (FDR < 0.01).

**Supplementary Table 24**| reactobiome consuming carbohydrates (i.e. mon-, di-, oligo- and poly saccharides) connected with plasma amino acids (FDR < 0.01). it shows the reactions along with corresponding sugar and sugar group as well as corresponding coefficient, p-value, and FDR of significantly associated plasma metabolites.

**Supplementary Table 25**| List of metabolites in biomass.

**Supplementary Table 26**| 153 pathways with KEGG reaction ID.

## References

1 Arumugam, M. et al. Enterotypes of the human gut microbiome. Nature 473, 174–180, doi:10.1038/nature09944 (2011).

2 Costea, P. I. et al. Enterotypes in the landscape of gut microbial community composition. Nat Microbiol 3, 8–16, doi:10.1038/s41564-017-0072-8 (2018).

3 Le Chatelier, E. et al. Richness of human gut microbiome correlates with metabolic markers. Nature 500, 541–546, doi:10.1038/nature12506 (2013).

4 Koh, A. & Backhed, F. From Association to Causality: the Role of the Gut Microbiota and Its Functional Products on Host Metabolism. Mol Cell 78, 584–596, doi:10.1016/j.molcel.2020.03.005 (2020).

5 Wilmanski, T. et al. Blood metabolome predicts gut microbiome alpha-diversity in humans. Nat Biotechnol 37, 1217–1228, doi:10.1038/s41587-019-0233-9 (2019).

6 Zierer, J. et al. The fecal metabolome as a functional readout of the gut microbiome. Nat Genet 50, 790–795, doi:10.1038/s41588-018-0135-7 (2018).

7 O’Brien, E. J., Monk, J. M. & Palsson, B. O. Using Genome-scale Models to Predict Biological Capabilities. Cell 161, 971–987, doi:10.1016/j.cell.2015.05.019 (2015).

8 Bordbar, A., Monk, J. M., King, Z. A. & Palsson, B. O. Constraint-based models predict metabolic and associated cellular functions. Nat Rev Genet 15, 107–120, doi:10.1038/nrg3643 (2014).

9 Tramontano, M. et al. Nutritional preferences of human gut bacteria reveal their metabolic idiosyncrasies. Nat Microbiol 3, 514–522, doi:10.1038/s41564-018-0123-9 (2018).

10 Magnusdottir, S. et al. Generation of genome-scale metabolic reconstructions for 773 members of the human gut microbiota. Nat Biotechnol 35, 81–89, doi:10.1038/nbt.3703 (2017).

11 Shoaie, S. et al. Quantifying Diet-Induced Metabolic Changes of the Human Gut Microbiome. Cell Metab 22, 320–331, doi:10.1016/j.cmet.2015.07.001 (2015).

12 Machado, D., Andrejev, S., Tramontano, M. & Patil, K. R. Fast automated reconstruction of genome-scale metabolic models for microbial species and communities. Nucleic Acids Res 46, 7542–7553, doi:10.1093/nar/gky537 (2018).

13 Norsigian, C. J., Fang, X., Seif, Y., Monk, J. M. & Palsson, B. O. A workflow for generating multi-strain genome-scale metabolic models of prokaryotes. Nat Protoc 15, 1–14, doi:10.1038/s41596-019-0254-3 (2020).

14 Diener, C., Gibbons, S. M. & Resendis-Antonio, O. MICOM: Metagenome-Scale Modeling To Infer Metabolic Interactions in the Gut Microbiota. mSystems 5, doi:10.1128/mSystems.00606-19 (2020).

15 Baldini, F. et al. The Microbiome Modeling Toolbox: from microbial interactions to personalized microbial communities. Bioinformatics 35, 2332–2334, doi:10.1093/bioinformatics/bty941 (2019).

16 Li, J. et al. An integrated catalog of reference genes in the human gut microbiome. Nat Biotechnol 32, 834–841, doi:10.1038/nbt.2942 (2014).

17 Ma, B. et al. A comprehensive non-redundant gene catalog reveals extensive within-community intraspecies diversity in the human vagina. Nat Commun 11, 940, doi:10.1038/s41467-020-14677-3 (2020).

18 Plaza Onate, F. et al. MSPminer: abundance-based reconstitution of microbial pan-genomes from shotgun metagenomic data. Bioinformatics 35, 1544–1552, doi:10.1093/bioinformatics/bty830 (2019).

19 Wu, Y. W., Simmons, B. A. & Singer, S. W. MaxBin 2.0: an automated binning algorithm to recover genomes from multiple metagenomic datasets. Bioinformatics 32, 605–607, doi:10.1093/bioinformatics/btv638 (2016).

20 Tebani, A. et al. Integration of molecular profiles in a longitudinal wellness profiling cohort. Nat Commun 11, 4487, doi:10.1038/s41467-020-18148-7 (2020).

21 Wen, C. et al. Quantitative metagenomics reveals unique gut microbiome biomarkers in ankylosing spondylitis. Genome Biol 18, 142, doi:10.1186/s13059-017-1271-6 (2017).

22 Norsigian, C. J. et al. BiGG Models 2020: multi-strain genome-scale models and expansion across the phylogenetic tree. Nucleic Acids Res 48, D402–D406, doi:10.1093/nar/gkz1054 (2020).

23 Holmes, I., Harris, K. & Quince, C. Dirichlet multinomial mixtures: generative models for microbial metagenomics. PLoS One 7, e30126, doi:10.1371/journal.pone.0030126 (2012).

24 Pedersen, H. K. et al. Human gut microbes impact host serum metabolome and insulin sensitivity. Nature 535, 376–381, doi:10.1038/nature18646 (2016).

25 Cotillard, A. et al. Dietary intervention impact on gut microbial gene richness. Nature 500, 585–588, doi:10.1038/nature12480 (2013).

26 Poole, J., Day, C. J., von Itzstein, M., Paton, J. C. & Jennings, M. P. Glycointeractions in bacterial pathogenesis. Nat Rev Microbiol 16, 440–452, doi:10.1038/s41579-018-0007-2 (2018).

27 Christgen, S. L. & Becker, D. F. Role of Proline in Pathogen and Host Interactions. Antioxid Redox Signal 30, 683–709, doi:10.1089/ars.2017.7335 (2019).

28 Gough, N. R. Proline Promotes Virulence. Science Signaling 3, ec31–ec31, doi:10.1126/scisignal.3106ec31 (2010).

29 Edwards, L. A. et al. Enterotoxin-producing staphylococci cause intestinal inflammation by a combination of direct epithelial cytopathy and superantigen-mediated T-cell activation. Inflamm Bowel Dis 18, 624–640, doi:10.1002/ibd.21852 (2012).

30 Ma, Y. et al. Seasonal variation in food intake, physical activity, and body weight in a predominantly overweight population. Eur J Clin Nutr 60, 519–528, doi:10.1038/sj.ejcn.1602346 (2006).

31 Elevated cardiac troponin T is an early indicator of anthracycline-induced cardiotoxicity. Nature Clinical Practice Oncology 2, 383–383, doi:10.1038/ncponc0237 (2005).

32 Gevers, D. et al. The treatment-naive microbiome in new-onset Crohn’s disease. Cell Host Microbe 15, 382–392, doi:10.1016/j.chom.2014.02.005 (2014).

33 Arany, Z. & Neinast, M. Branched Chain Amino Acids in Metabolic Disease. Curr Diab Rep 18, 76, doi:10.1007/s11892-018-1048-7 (2018).

34 Mika, A. et al. A comprehensive study of serum odd- and branched-chain fatty acids in patients with excess weight. Obesity (Silver Spring) 24, 1669–1676, doi:10.1002/oby.21560 (2016).

35 Wang, R. et al. The prevalence of pre-diabetes and diabetes and their associated factors in Northeast China: a cross-sectional study. Sci Rep 9, 2513, doi:10.1038/s41598-019-39221-2 (2019).

36 Hu, C. & Jia, W. Diabetes in China: Epidemiology and Genetic Risk Factors and Their Clinical Utility in Personalized Medication. Diabetes 67, 3–11, doi:10.2337/dbi17-0013 (2018).

37 Warnatsch, A. et al. Reactive Oxygen Species Localization Programs Inflammation to Clear Microbes of Different Size. Immunity 46, 421–432, doi:10.1016/j.immuni.2017.02.013 (2017).

38 Yoo, S. K. & Huttenlocher, A. Innate immunity: wounds burst H2O2 signals to leukocytes. Curr Biol 19, R553–555, doi:10.1016/j.cub.2009.06.025 (2009).

39 de Castro, J. M. Seasonal rhythms of human nutrient intake and meal pattern. Physiol Behav 50, 243–248, doi:10.1016/0031-9384(91)90527-u (1991).

40 Cronise, R. J., Sinclair, D. A. & Bremer, A. A. The “metabolic winter” hypothesis: a cause of the current epidemics of obesity and cardiometabolic disease. Metab Syndr Relat Disord 12, 355–361, doi:10.1089/met.2014.0027 (2014).

41 Rinke, C. et al. Insights into the phylogeny and coding potential of microbial dark matter. Nature 499, 431–437, doi:10.1038/nature12352 (2013).

42 Wheeler, T. J. & Eddy, S. R. nhmmer: DNA homology search with profile HMMs. Bioinformatics 29, 2487–2489, doi:10.1093/bioinformatics/btt403 (2013).

43 Seaver, S. M. D. et al. The ModelSEED Biochemistry Database for the integration of metabolic annotations and the reconstruction, comparison and analysis of metabolic models for plants, fungi and microbes. Nucleic Acids Res, doi:10.1093/nar/gkaa1143 (2020).

44 Arkin, A. P. et al. KBase: The United States Department of Energy Systems Biology Knowledgebase. Nat Biotechnol 36, 566–569, doi:10.1038/nbt.4163 (2018).

45 Heirendt, L. et al. Creation and analysis of biochemical constraint-based models using the COBRA Toolbox v.3.0. Nat Protoc 14, 639–702, doi:10.1038/s41596-018-0098-2 (2019).

46 Wang, H. et al. RAVEN 2.0: A versatile toolbox for metabolic network reconstruction and a case study on Streptomyces coelicolor. PLoS Comput Biol 14, e1006541, doi:10.1371/journal.pcbi.1006541 (2018).

47 Masood, W., Annamaraju, P. & Uppaluri, K. R. in StatPearls (2020).

48 Te Morenga, L. A., Levers, M. T., Williams, S. M., Brown, R. C. & Mann, J. Comparison of high protein and high fiber weight-loss diets in women with risk factors for the metabolic syndrome: a randomized trial. Nutr J 10, 40, doi:10.1186/1475-2891-10-40 (2011).

49 Gu, Z., Gu, L., Eils, R., Schlesner, M. & Brors, B. circlize Implements and enhances circular visualization in R. Bioinformatics 30, 2811–2812, doi:10.1093/bioinformatics/btu393 (2014).

50 Shannon, P. et al. Cytoscape: a software environment for integrated models of biomolecular interaction networks. Genome Res 13, 2498–2504, doi:10.1101/gr.1239303 (2003).

51 Johansson, M. E., Larsson, J. M. & Hansson, G. C. The two mucus layers of colon are organized by the MUC2 mucin, whereas the outer layer is a legislator of host-microbial interactions. Proc Natl Acad Sci U S A 108 Suppl 1, 4659–4665, doi:10.1073/pnas.1006451107 (2011).

52 Henry, C. S. et al. High-throughput generation, optimization and analysis of genome-scale metabolic models. Nat Biotechnol 28, 977–982, doi:10.1038/nbt.1672 (2010).

53 Wagner, G. P., Kin, K. & Lynch, V. J. Measurement of mRNA abundance using RNA-seq data: RPKM measure is inconsistent among samples. Theory Biosci 131, 281–285, doi:10.1007/s12064-012-0162-3 (2012).

